# Reproducible Risk Loci and Psychiatric Comorbidities in Anxiety: Results from ^~^200,000 Million Veteran Program Participants

**DOI:** 10.1101/540245

**Authors:** Daniel F. Levey, Joel Gelernter, Renato Polimanti, Hang Zhou, Zhongshan Cheng, Mihaela Aslan, Rachel Quaden, John Concato, Krishnan Radhakrishnan, Julien Bryois, Patrick F. Sullivan, Million Veteran Program, Murray B. Stein

## Abstract

Anxiety disorders are common and often disabling. They are also frequently co-morbid with other mental disorders such as major depressive disorder (MDD); these disorders may share commonalities in their underlying genetic architecture. Using one of the largest homogenously phenotyped cohorts available, the Million Veteran Program sample, we investigated common variants associated with anxiety in genome-wide association studies (GWASes), using survey results from the GAD-2 anxiety scale (as a continuous trait, n=199,611), and self-reported anxiety disorder diagnosis (as a binary trait, n=224,330). This largest GWAS to date for anxiety and related traits identified numerous novel significant associations, several of which are replicated in other datasets, and allows inference of underlying biology.

**Abstract:** We used GWAS in the Million Veteran Program sample (nearly 200,000 informative individuals) using a continuous trait for anxiety (GAD-2) to identify 5 genome-wide significant (GWS) signals for European Americans (EA) and 1 for African Americans. The strongest findings were on chromosome 3 (rs4603973, p=7.40×10^−11^) near the *SATB1* locus, a global regulator of gene expression and on chromosome 6 (rs6557168, p=1.04×10^−9^) near *ESR1* which encodes estrogen receptor α. A locus identified on chromosome 7 near *MADIL1* (p=1.62×10^−8^) has been previously identified in GWAS of bipolar disorder and of schizophrenia and may represent a risk factor for psychiatric disorders broadly. SNP-based heritability was estimated to be ~6% for GAD-2. We also GWASed for self-reported anxiety disorder diagnoses (N=224,330) and identified two GWS loci, one (rs35546597, MAF=0.42, p=1.88×10^−8^) near the *AURKB* locus, and the other (rs10534613, MAF=0.41, p=4.92×10^−8^) near the *IQCHE* and *MADIL1* locus identified in the GAD-2 analysis. We demonstrate reproducibility by replicating our top findings in the summary statistics from the Anxiety NeuroGenetics Study (ANGST) and a UK Biobank neuroticism GWAS. We also replicated top findings from a large UK Biobank preprint, demonstrating stability of GWAS findings in complex traits once sufficient power is attained. Finally, we found evidence of significant genetic overlap between anxiety and major depression using polygenic risk scores, but also found that the main anxiety signals are independent of those for MDD. This work presents novel insights into the neurobiological risk underpinning anxiety and related psychiatric disorders.

## Introduction

Anxiety disorders are common, affecting 1 in 10 Americans each year^1^, and are a leading cause of disability worldwide.^2^ A recent analysis of anxiety and depressive disorders showed that cumulatively they cost about $90 billion in personal health spending for 2013.^3^ Given their prevalence, associated impairment, and economic costs, anxiety disorders are a major public health concern.^4^

Anxiety is “a future-oriented mood state associated with preparation for possible, upcoming negative events”,^5^ and is usually a normal and adaptive behavioral response to everyday life. In anxiety *disorders*, anxiety is excessive or out of proportion to the actual or anticipated event, and accompanied by clinically significant distress or disability.^6^ Numerous risk factors for anxiety disorders have been studied, including experiential and genetic factors.^7^ For example, neurotic personality traits are predictive of the onset of anxiety disorders.^8^ Twin studies demonstrate a heritable component to anxiety disorders,^7^ but there have been few published GWAS to date investigating anxiety or anxiety related traits. The Anxiety Neuro Genetics Study (ANGST)^9^ meta-analysis has been the most noteworthy published study to date, finding one genome wide significant (GWS) locus each for a categorical case control design for any anxiety disorder diagnosis and a quantitative factor score for anxiety in a cohort of over 18,000 subjects. Understanding of the genetics of anxiety disorders has thus lagged behind other related disorders such as major depression.^10^

Only a third of individuals with anxiety disorders receive treatment.^11^ For those who do enter treatment, psychological approaches such as cognitive behavioral therapy have been shown to be very effective^12;13^; as have certain pharmacotherapies.^14,15^, Better understanding of genetic risk factors and determinants now inform other aspects of medicine such as oncology and cardiology through identification of causal mutations^16^ and variants, and will have important implications for psychiatry.^17^ These precision medicine approaches are challenging in complex traits such as anxiety because they are associated with many (perhaps hundreds of thousands) of variants of individually small effect^18^. The use of polygenic risk scores will require a suitably large sample size to provide sufficient confidence in these small individual effects that cumulatively account for so much of the heritability. Underlying polygenic risk factors from sufficiently large cohorts may inform an approach to identify those with a predisposition to develop anxiety disorders and improve outcomes.

The Million Veteran Program (MVP) is one of the world’s largest databases of genetic, environmental, and medical information, based on data from United States military veterans^19^. Using this large genetic dataset and (1) the Generalized Anxiety Disorder 2-item (GAD-2) scale^20^ as well as (2) self-report of physician diagnosis of anxiety or panic disorder, we discovered novel GWS variants associated with anxiety in European Americans (EA) and African Americans (AA). We examined replication and genetic overlap of these results with those of previous studies examining anxiety and traits with which anxiety disorders are commonly comorbid – major depression, PTSD, and neuroticism. We also examined expression quantitative trait loci (eQTLs) to identify possible gene expression implications of these genetic variants, with eQTL evidence for altered expression in the basal ganglia and cerebellum. These findings, in the largest cohort of individuals GWASed for anxiety and anxiety disorders (199,611 subjects for the quantitative trait, 224,330 for binary diagnosis), indicate shared genetic risk with some other mental disorders, but also point to loci that may be especially important for anxiety and anxiety-related traits.

## Methods

### Participants

The MVP cohort has been described previously.^19^ Results were analyzed in two separate tranches based on when genotyping results were conducted. Ancestry was assigned using 10 principal components and the 1000 Genomes project phase 3 EUR and AFR as reference within each tranche of data.

### Genotyping, Imputation, and Quality Control

Genotyping, imputation, and quality control within MVP has been previously described. Briefly, samples were genotyped using a 723,305-SNP Affymetrix Axiom biobank array, customized for MVP.^19^ Imputation was performed with minimac3 using data from the 1000 Genomes project. For post-imputation QC, SNPs with imputation INFO scores of < 0.3 or minor allele frequencies (MAF) below 0.01 were removed from analysis. For the first tranche of data, 22,183 SNPs were selected through linkage disequilibrium (LD) pruning using PLINK,^21^ and then Eigensoft^22^ was used to conduct principal component analysis on 343,286 MVP samples and 2,504 1000 Genomes samples.^23^ The reference population groups (EUR, EAS, AFR, AMR, or SAS) in the 1000 Genomes samples were used to define EUR (n=241,541) and AFR (n=61,796) groups used in this analysis. Similar methods were used in the second tranche of data, which contained 108,416 new MVP samples and the same 2,504 1000 Genomes samples. In Tranche 2, 80,694 participants were defined as EUR and 20,584 were defined as AFR.

### Phenotypic Assessment

The GAD-2 scale consists of 2 questions (Table S1) included in a self-report survey, each scored on a 0-3 scale. Participants were asked to respond based on their symptoms during the past 2 weeks. Values for the two responses are summed, resulting in a range of scores between 0-6, which we treated as a continuous trait (Table 1).

Another anxiety phenotype was analyzed based on data collected from the MVP baseline survey. Participants were asked the question: “Please tell us if you have been diagnosed with the following conditions: Anxiety reaction/Panic disorder.” Answers were recorded as yes/no binary responses and missing responses were excluded from analysis. 224,330 participants (34,189 cases) responded to the diagnosis question and, had available genotype, and had either a European or African ancestry assigned.

### Statistical Analysis

GWAS analysis was carried out by linear regression for each ancestry group and tranche using PLINK 2.0 on dosage data, covarying for age, sex, and the first 10 PCs. Meta-analysis was performed using METAL (EA n=175,163, AA n=24,448). Similar analyses were conducted for the selfreported medical history of anxiety diagnosis except that logistic regression was performed.

### LD Score Regression (LDSC) and SNP-based Heritabilitv

We used LD score regression (LDSC) through LD Hub^24^ to estimate SNP-based heritability, and to assess genetic correlation of GAD-2 anxiety with 35 traits selected using the category keywords: psychiatric, personality, cognitive, reproductive, sleeping, smoking behavior, aging, and neurological. The traits from the ANGST GWAS of anxiety case-control and factor scores^9^ were calculated separately in LDSC using summary statistics downloaded from the Psychiatric Genomics Consortium website.

### Conditional Analysis for Major Depression

Considering the extensive comorbidity between major depression and anxiety disorders^7^ – we ran a conditional analysis with the mtCOJO method^25^ using GCTA software with the MVP GAD-2 summary statistics as the primary analysis and the PGC MDD2 (excluding 23andMe)^10^ summary statistics to condition the analysis for depression.

### Gene-based Tests

Summary statistics from the GWAS were loaded into Functional Mapping and Annotation of Genome-Wide Association Studies (FUMA GWAS) to test for gene-level associations using Multi-Marker Analysis of GenoMic Annotation (MAGMA).^26,27^ Input SNPs were mapped to 17,927 protein coding genes. The GWS threshold for the gene-based test was therefore determined to be p = 0.05/17,927 = 2.79×10^−6^.

## Results

### Primary analysis

GWAS was conducted separately in two tranches of each ancestry in the MVP sample, defined by the time data became available, and meta-analyzed together within population. One genomic locus was GWS in the AA meta-analysis (Figure 1a) while five loci were GWS in the EA metaanalysis (Figure 1b).

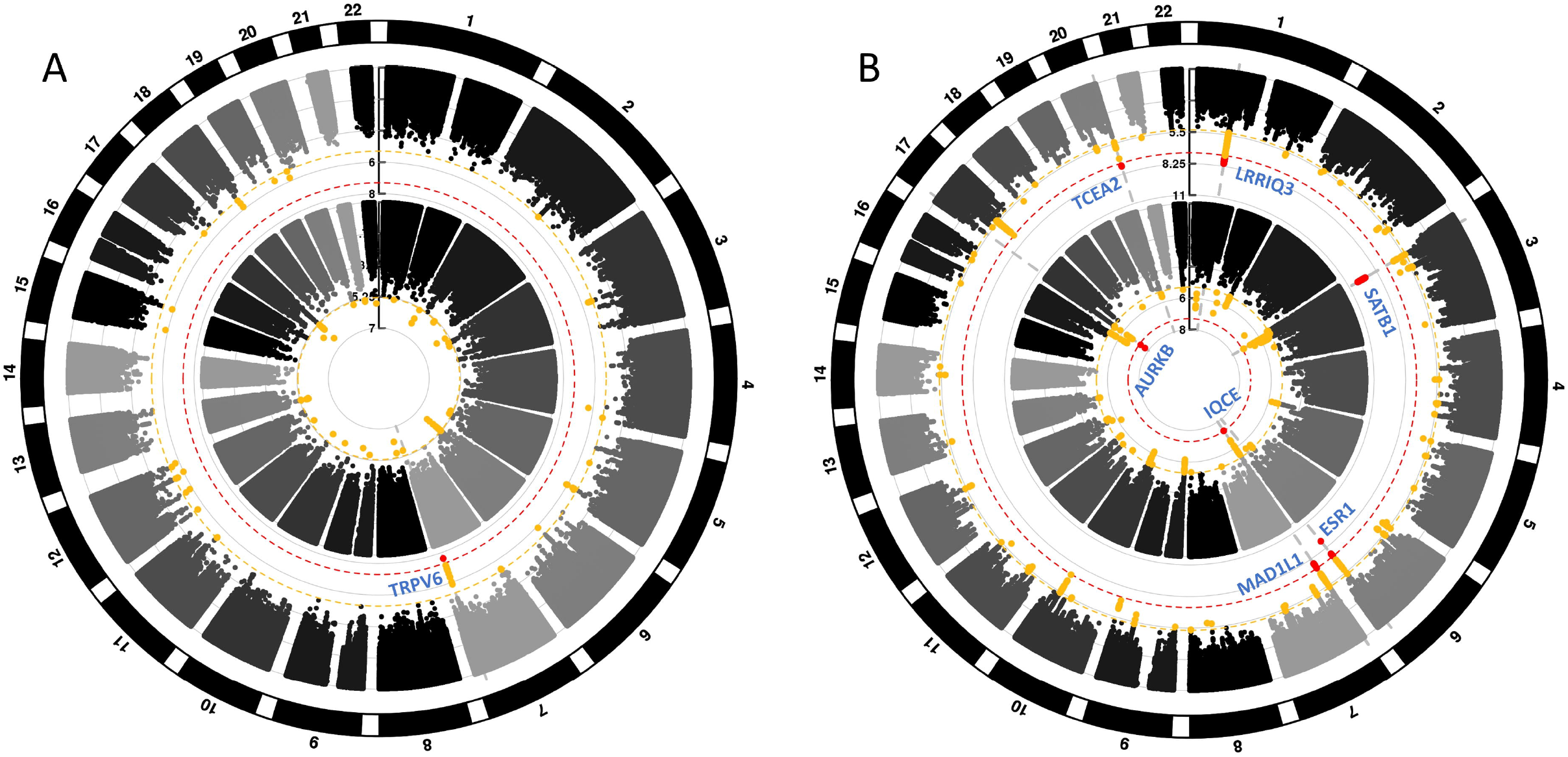

The GWS result from the AA analysis was near the *TRPV6* (Transient Receptor Potential Cation Channel Subfamily V Member 6) locus. The top signal in the EA meta-analysis consisted of 64 GWS SNPs in high LD at the *SATB1-AS1* (Special AT-Rich Sequence Binding 1 Antisense RNA 1) locus. The strongest finding (rs4603973, MAF=0.29, p=7.40×10^−11^) was intronic at *SATB1-AS1*. The second strongest independent signal was on chromosome 6 (rs6557168, MAF=0.63, p=1.04×10”^9^) intronic at *ESR1* (Estrogen Receptor 1) with 10 other GWS SNPs in high LD. A third GWS association for EAs was found on chromosome 1 (rs12023347, MAF=0.48, p=9.73×10^−9^) near the long noncoding RNA *LINC01360* and *LRRIQ3*. The fourth GWS association found in EAs was on chromosome 7 (rs56226325, MAF=0.17, p=1.62×10^−8^) in an intron of *MAD1L1* (Mitotic Arrest Deficient 1 Like 1). The fifth association for EAs was in and around the *TCEA2* (Transcription Elongation Factor A2), *RGS19* (Regulator Of G Protein Signaling 19), and *OPRL1* (Opioid Related Nociceptin Receptor 1) genes (rs6090040, MAF=0.48, p=2.59×10^−8^).

We conducted additional analysis in related phenotypes within the same samples using case-control status for self-reported diagnosis of anxiety or panic disorder. For the EA subjects, there were two GWS signals for the self-reported Anxiety/Panic case control GWAS, in a gene-rich region nearest *AURKB* on chromosome 17 (rs35546597, MAF=0.42, p=1.88×10^−8^) and chromosome 7 in an *IQCHE* intron (rs10534613, MAF=0.41, p=4.92×10^−8^) close to the *MAD1L1* locus identified for GAD-2. There were no GWS findings for this phenotype in African Americans.

### Replication

For replication we looked up our top 5 SNPs in 3 independent GWAS with anxiety-related phenotypes. We looked up our lead GWS SNPs in GWAS for anxiety case-control,^9^ and neuroticism in the recent large UK Biobank, 23andMe, and Genetics of Personality Consortium^28^ (Table 2). (The lead SNP on chromosome 3 near the *SATB1* locus, rs4603973, was not available for lookup in the UK Biobank neuroticism GWAS, so we used the strongest LD-proxy available (rs4390955 R^2^=0.91, p=8.68E-11), as a surrogate.) In the UK Biobank neuroticism study, 4 of 5 independent SNPs looked up had the same direction of effect, 3 were significant, and one near MADIL1 was nearly GWS (rs56226325, p=6.59×10-8). A preprint reported results for anxiety from UK Biobank using Case-Control and the GAD-7, scored as a dichotomous trait.^29^ We find significant replication for 2 of their 4 findings, with suggestive evidence for a third (Table 3).

### Genome wide gene based association study (GWGASD for GAD-2

In the GWGAS, the top gene identified was *OPRL1* (p=1.15×10^−9^), which was also significant in the SNPwise analysis as noted above. 31 genes were identified as GWS following Bonferroni correction for multiple comparisons. A more permissive Benjamini-Hochberg correction with a still relatively restrictive 0.05 FDR identified 189 genes (Supplementary Table S3) in total for investigation of biological relevance through the Ingenuity pathway enrichment tool. Ingenuity Pathway Analysis (IPA, Ingenuity Systems, www.ingenuity.com, Redwood City, CA, USA) was used to examine underlying biological process for the 189 genes corrected by Benjamini-Hochberg (Supplementary Table S4).

### eQTL

To identify causal implications for genetic variants, eQTLs were assessed for top GWS signals using GTEx v7 brain tissue expression data. Top GWS signals on chromosome 7 and 20 had significant eQTL (FDR <= 0.05) for 4 different genes: *FTSJ2, RGS19, C20orf201*, and *OPRL1* (Supplementary table S7). The top signals are centered in the basal ganglia and cerebellum.

### LD score regression (LDSC) analysis

Eighteen of the 35 traits were significantly correlated with GAD-2 following correction for multiple comparisons (α= 0.05/35= 0.001429) (Figure 2 and Supplementary Table S5). The most significantly correlated traits were depressive symptoms (r_g_=0.8055, p=1.95×10^−53^) and neuroticism (r_g_=0.7174, p=6.53×10^−53^).

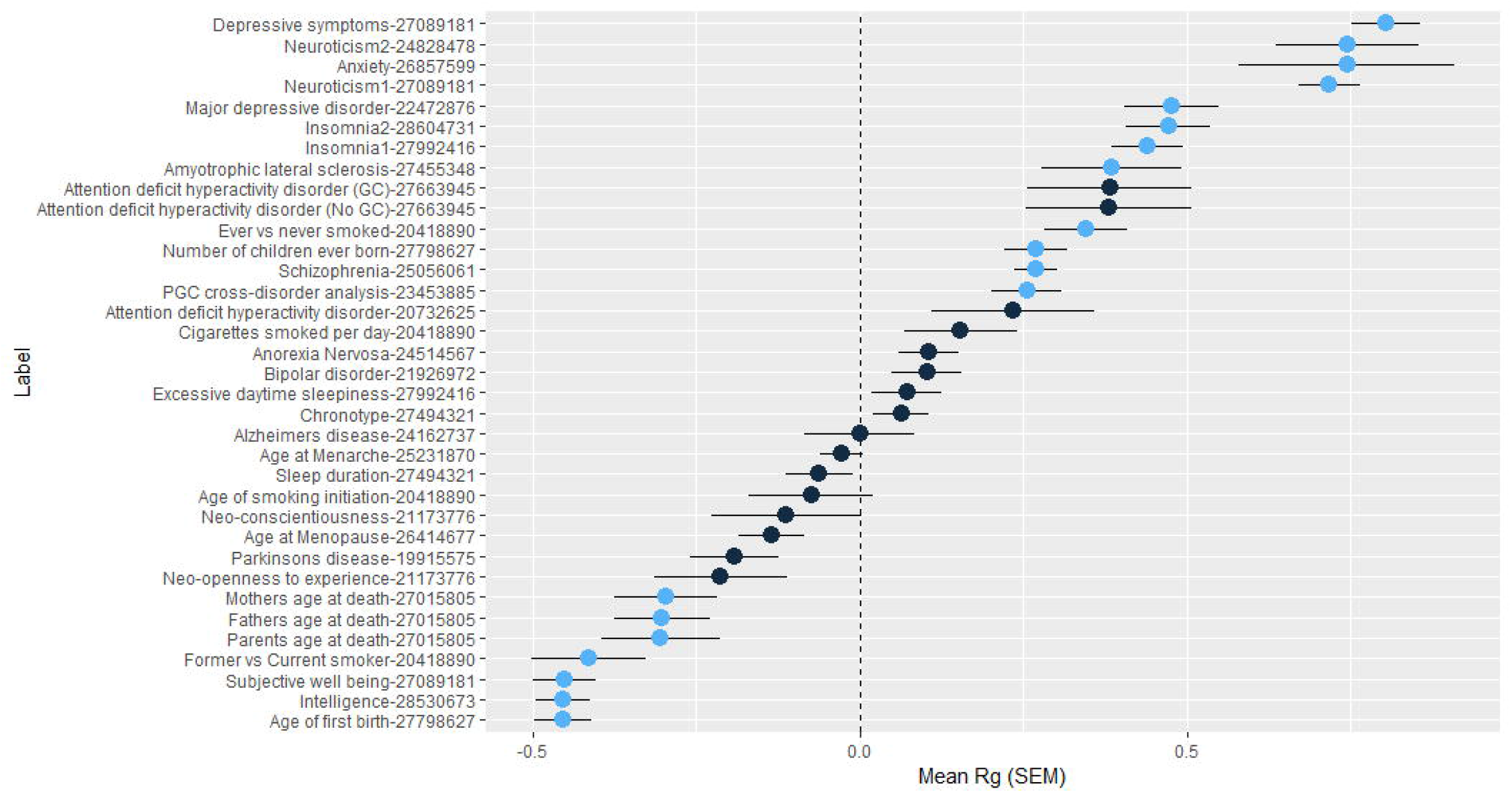

We also investigated genetic correlation within the MVP cohort using the GWAS for selfreported case-control anxiety. Genetic correlations between GAD-2 and self-reported diagnosis were high, with an r_g_ of 0.856. The phenotype correlation between GAD-2 and self-reported diagnosis was r=0.64, p<2.2×10^−16^.

### SNP heritability

SNP-based heritability using LDSC for the GAD-2 quantitative trait was estimated to be 0.0558 (SE = 0.004). SNP based inflation was mild considering the sample size and polygenic trait studied (λ =1.19); the intercept (1.026) and attenuation ratio (0.1177) estimated by LDSC showed negligible evidence for inflation due to population stratification. SNP based heritability for the phenotype of case-control anxiety was 0.0374 (SE= 0.004).

### Polygenic risk score (PRS) analysis

was used to test polygenic risk scores. Summary statistics from the MVP GAD-2 analysis were used as the base data for calculating the PRS (using PRSice v 1.25^30^). Genetic overlap between anxiety and PTSD or MDD were tested in the Psychiatric Genomics Consortium (PGC) MDD^10^ or PTSD^31^ GWAS, respectively, and overlap with case-control anxiety disorder was tested in the largest previously available study.^9^ Significant overlap was identified: the MVP GAD-2 PRS can explain up to 0.24% of the variance in MDD in the PGC GWAS (p=2.05×10^−94^), 0.23% of the variance of PTSD in the PGC GWAS (p=4.23×10^−12^), and 0.48% of the variance in ANGST Anxiety Disorder (p=3.66×10^−20^) (Figure 3).

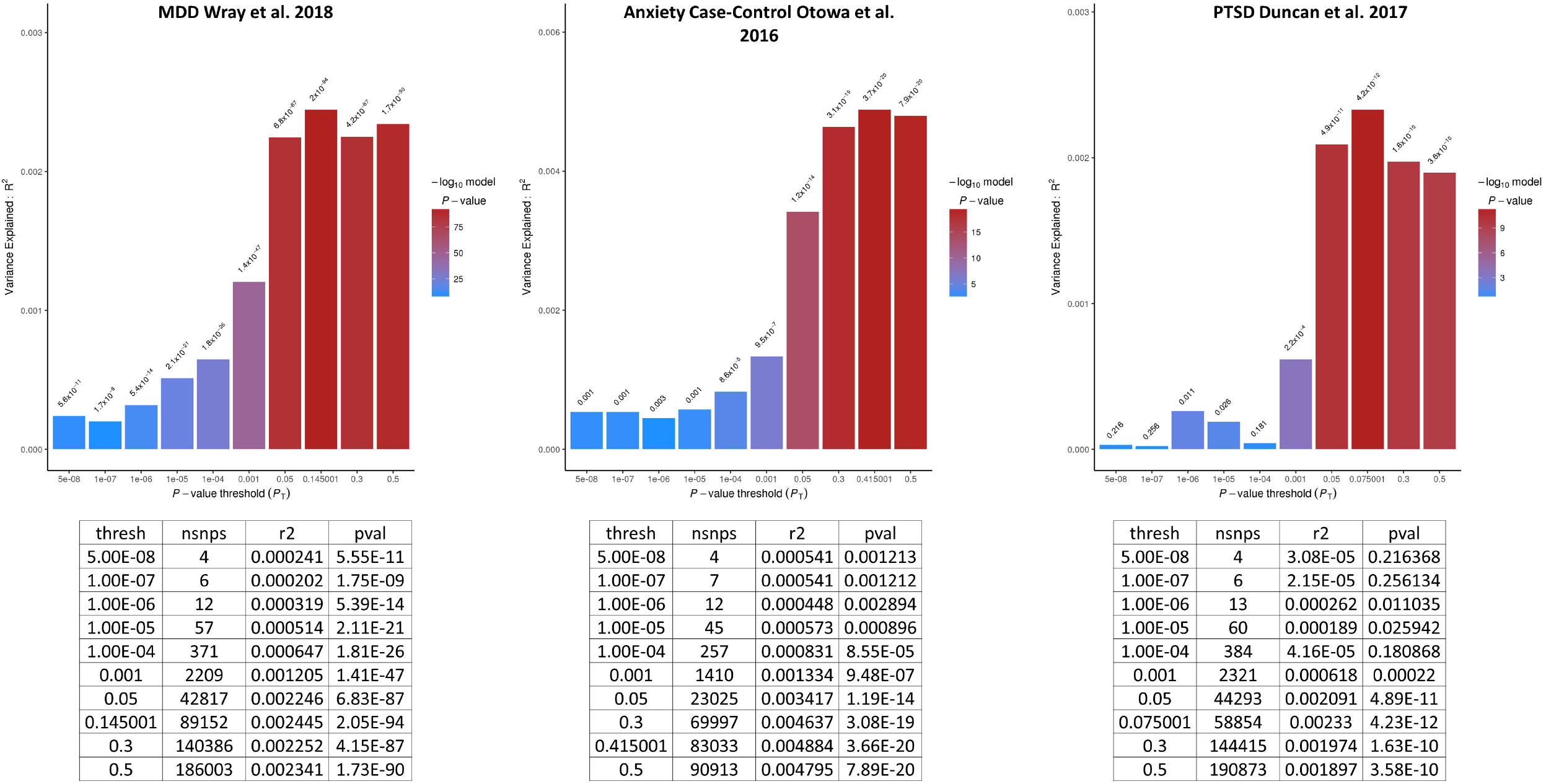

### mtCOJO

mtCOJO GWAS was used to condition the GAD-2 MVP summary statistics for anxiety on the PGC MDD2 summary statistics for MDD^10^. There were no new signals and significance of lead findings dropped by about an order of magnitude, but results on chromosomes 3 (*SATB1*) and 6 *(ESR1)* remained GWS.

## Discussion

We present the largest GWAS to date for anxiety traits, employing a quantitative phenotype, the GAD-2, in nearly 200,000 MVP subjects, as well as self-reported anxiety/panic case-control phenotypes in >220,000 MVP subjects. We identified novel genetic variants in and around genes some of which have previously-known functional relationships with anxiety. These genes play roles in the HPA axis, neuronal development, and global regulation of gene expression

There is high comorbidity between anxiety and PTSD.^32^ We used a PRS derived from the MVP GAD-2 analysis to identify genetic overlap with the independent PGC PTSD GWAS (Figure 3). We found a high degree of genetic overlap between these two traits, providing biological evidence that this known clinical comorbidity is due at least in part to shared genetic etiology. We also used LDSC to identify correlation between traits relevant to brain and behavior. We identified substantial positive correlations with depression and neuroticism as well as a negative correlation with subjective well-being (Figure 2).

The GWS result in AAs is an insertion variant which is rare outside of African ancestry and occurs in a genomic region proposed to be under recent selection in Europeans.^33^ The lead SNP is at *TRPV6*, which encodes a Ca2+ selective^34^ membrane cation channel associated with epithelial calcium transport and homeostasis in kidney and intestine. The lead SNP rs575403075 has an MAF range of between 0% and 1% in non-African populations and would often fall below MAF QC thresholds used for common variants in most non-African populations. This highlights the importance of studying genetics in diverse populations, otherwise these signals may be missed entirely.

The top GWS findings for EAs in the GAD-2 analysis were in and around *SATB1* and the antisense gene *SATB1-AS1. SATB1* is a global regulator that influences expression of multiple genes^35^ involved in neuronal development. One gene modulated in expression is Corticotropin Releasing Hormone (*CRH*), encoding the protein product of the same name which plays an essential role in the HPA axis, which has frequently been shown to modulate stress and fear/anxiety response.^36^ The *CRHR1* (Corticotropin Releasing Hormone Receptor 1) gene was GWS in the gene-based association analysis (p=3.60×10^−7^). CRHR1 is bound by endogenous CRH and CRHR1 has been a proposed target for treatment, with evidence for anxiolytic-like effects of CRHR1 antagonists in animal models^36,37^ although not yet in humans.^38,39^ One possible explanation for a lack of anxiolytic effect observed in humans is heterogeneity in the patient population. The present results suggest that this may be a reasonable pathway to address via personalized medicine, as individuals with differing genetic risk that does or does not involve this pathway may differ in their responses to glucocorticoid-targeted therapeutic agents.

The estrogen receptor *ESR1* (also known as estrogen receptor α) has been a focus in animal models of anxiety-like behaviors and these have provided mechanistic validity for the role of ESR1. Studies of estradiol administration to ovariectomized rats and ESR1 null mice have shown consistent evidence that ESR1 is involved in anxiety-like behavior in non-human animals.^40^ Our finding of an association between *ESR1* and anxiety may have implications for our understanding of sex differences in anxiety disorders, which generally show about a 2:1 female predominance, and greater susceptibility to develop PTSD following traumatic events.^41^ Although this increased susceptibility is partially explained by sex-specific exposure to certain kinds of traumatic events, there may also be differential biological context provided in part by the role of the estrogen receptor. Our study in a predominantly male sample identifies *ESR1* as GWS. Estrogen is important in both sexes, with a recent review highlighting the important role for estrogens in men.^42^ Future studies with larger sample sizes of women should continue to investigate sex differences in genetic risk for anxiety-related traits.

The lead SNP from the GAD-2 GWAS near the *LINCO1360* and *LRRIQ3* (rs2180945) loci is nominally significant and has the same direction of effect in the PGC MDD 2018 analysis (p= 1.434×10^−6^). Previous genetic epidemiology studies have shown that common genetic factors can underlie anxiety and depressive traits.^43^ This variant may be linked to a common risk factor for both disorders.

One GWS signal for GAD-2 was in a gene-rich region on chromosome 20 near *TCEA2, C20orf201, RGS19*, and *OPRL1*. eQTL data suggest that variants in this region play roles in the regulation of the expression of *RGS19* and *OPRL1* in the cerebellum and in the basal ganglia (Supplementary Table S7). Increased *OPRL1* expression has been found in the amygdala of PTSD-like severe stress exposed mouse model following a cued-fear expression paradigm.^44^

*MADIL1* (GAD-2 lead SNP rs56226325, MAF=0.17, p=4.68×10^-10^, case-control lead SNP rs10534613, MAF=0.41, p=4.92×10^−8^) has been associated previously with bipolar disorder^45^ and one of the lead SNPs in that study in a *MAD1L1* intron is also nominally associated with anxiety in the current study (rs11764590, p= 2.82×^−7^). This locus has also been identified among 108 GWS loci by the PGC schizophrenia study,^46^ and our lead SNP is nominally significant in that study (rs56226325, p=1.12×10^−3^). This SNP is also nominally significant (6.71×10^−4^) in the 2018 PGC depression GWAS ^10^, so this locus may be a common risk factor for several psychiatric disorders.

*MADIL1\**rs56226325 is also an eQTL with the gene *FTSJ2* (Supplementary Table S7), which exhibits altered expression in the brain. This gene is a mitochondrial RNA methyltransferase important for the proper assembly of the mitochondrial ribosome and cellular respiration.^47^ The protein product of *FTSJ2* is “Mitochondrial rRNA Methyltransferase 2” (*MRM2*), which was implicated in a case-study of a 7 year old Italian boy with a damaging mutation leading to an encephalopathy, lactic acidosis and stoke-like (MELAS) syndrome.^48^ Larger effect mutations at this locus can have devastating effects on the brain; smaller effect variations may be less deleterious but still cumulatively influence development which may predispose to neurological and psychiatric disorders.

The Brainstorm Consortium has investigated shared heritability between psychiatric and neurological disorders.^49^ Consistent with their findings, we find very strong genetic correlation between anxiety and psychiatric traits such as depression (r_g_ = 0.81, p=1.95×10^−53^) and neuroticism (r_g_ = 0.72, p=6.53×10^−53^) but relatively weaker genetic overlap with neurological disorders. Significant overlap is detected for amyotrophic lateral sclerosis (r_g_ = 0.39, p=3.00×10^−4^) but Parkinson’s disease (r_g_ = −0.19, p=4.80×10^−3^) doesn’t survive correction for multiple comparisons and no overlap is seen for Alzheimer’s disease (r_g_ = 0.00, p=1.00). The relatively strong genetic overlap across psychiatric disorders is a common feature in psychiatric genetics and could point to a need for increased diagnostic specificity, perhaps with a focus on more dimensional rather than diagnostic phenotypes. This may also point to opportunities to improve treatment where there is evidence for greater biological and genetic overlap between disorders. The blurred boundaries between some psychiatric disorders in the clinic are increasingly supported by genetic findings.^10 31 49^

Limitations to this work include the fact that phenotypes were based on self-reported survey data. The GAD-2 asks questions which temporally reference the “past 2 weeks,” that is, it assesses state rather than trait anxiety. This measure should still capture a large proportion of those who experience anxiety on a daily or regular basis, but it falls short of the desired trait (lifetime) anxiety measure. Similarly, the question about diagnosis for anxiety or panic relies on self-report. Also, MVP has mostly male participants. While women are included in this analysis, clinically important interactions between sex, phenotype, and genotype could not be addressed. This cohort is growing, and future recruitment will provide additional power to revisit sex stratified analyses of this sample.

In summary, we have identified novel variants for anxiety by performing GWAS in the large MVP cohort. We found replication of many of these findings in independent cohorts assessed with similar or related phenotypes. We replicated results in our GWAS for top findings from other relevant trait-related GWAS. We also identified significant genetic overlap with MDD, PTSD, and Neuroticism using polygenic risk scores and LD score regression. This work provides additional genetic evidence for the overlap between disorders which frequently present co-morbid with anxiety and presents new molecular targets for investigation with a longer view toward the development of new treatments.

## Acknowledgements

The authors would like the thank the veterans who participate in the Million Veterans Program, without whom this work could not have been possible. This study was supported by funding from the Veteran Affairs Office of Research and Development Million Veteran Program Grant#MVP011 and CSP575B.

## Competing Interests

Dr. Stein reports receiving consulting fees in the past 3 years from Aptinyx, Bionomics, Janssen, Neurocrine, Pfizer, and Oxeia Biopharmaceuticals. All other authors declare that they have no conflict of interest.

