## Supplementary Material for "Reproducible Risk Loci and Psychiatric Comorbidities in Anxiety: Results from ^~^200,000 Million Veteran Program Participants"

Supplementary Figure S1

Manhattan Plot for EA Anxiety/Panic Case-Control GWAS

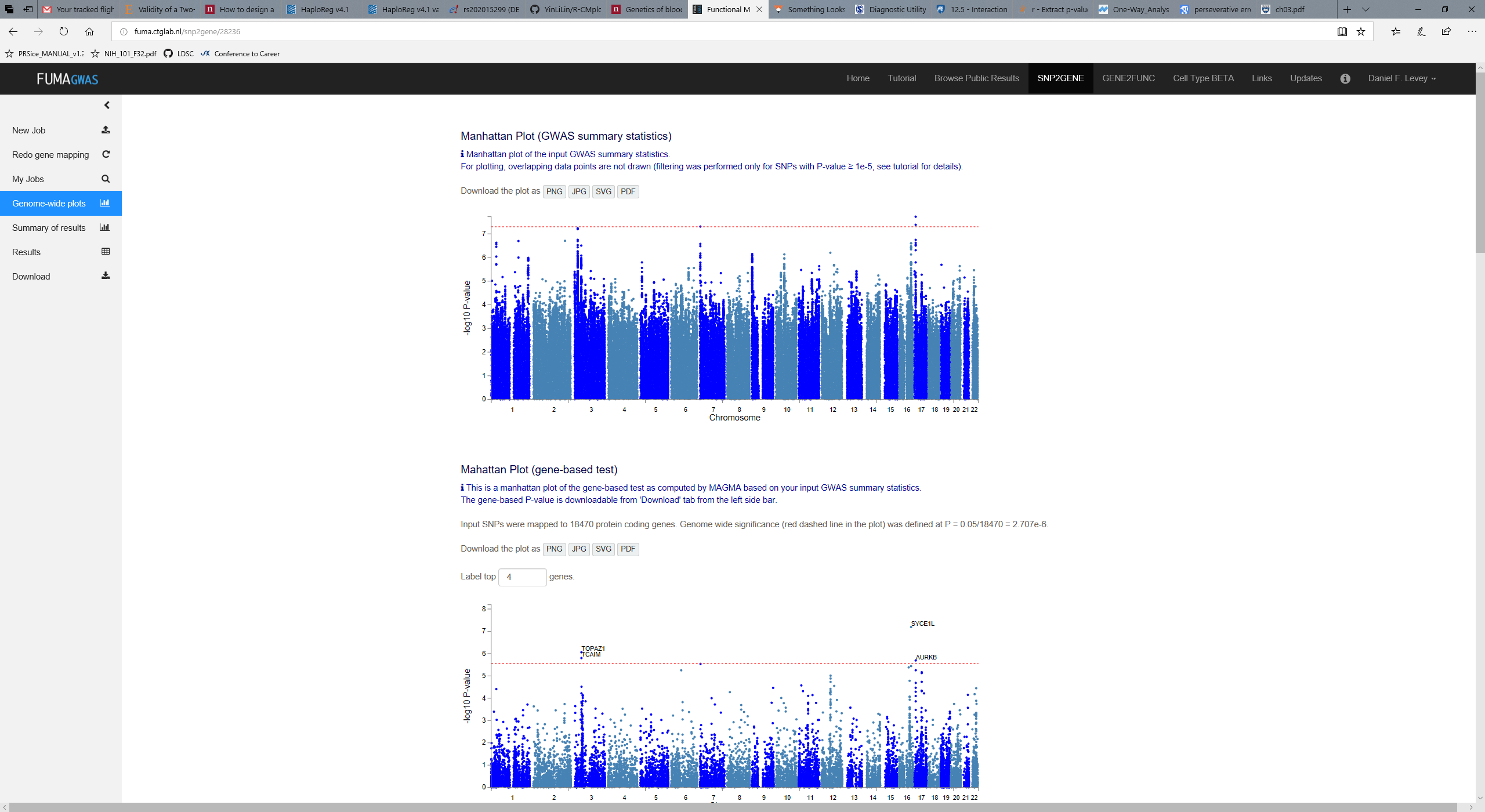

Supplementary Figure S2

Manhattan plot for AA Anxiety/Panic Case-Control GWAS

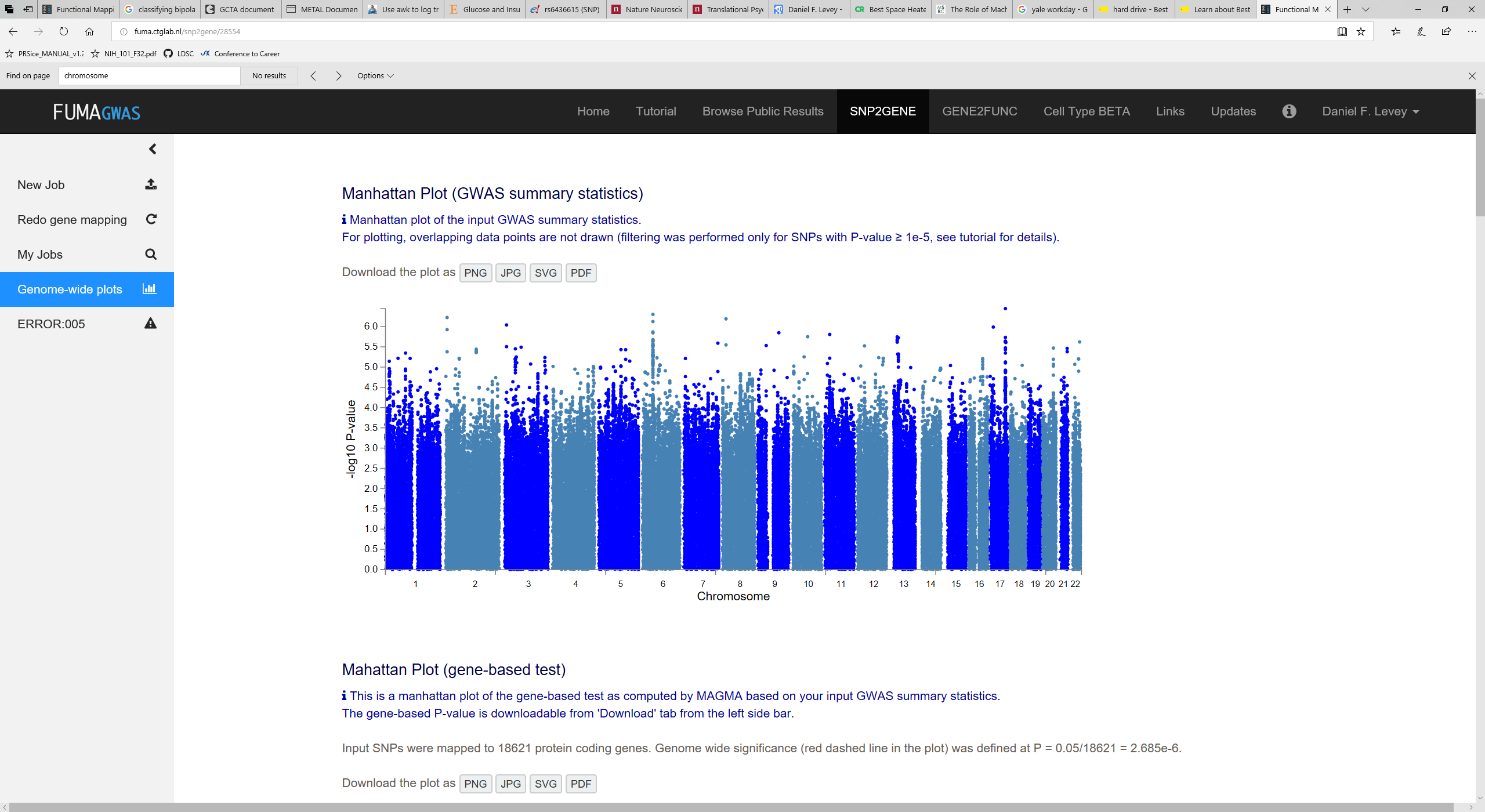

Supplementary Figure S3. Conditional Analysis with mtCOJO using the PGC MDD2 summary statistics (excluding 23andMe). Top is the original GWAS of GAD-2, bottom is the conditional analysis. In most cases signal was reduced by an order of magnitude; peaks on chromosomes 3 and 6 remain GWS.

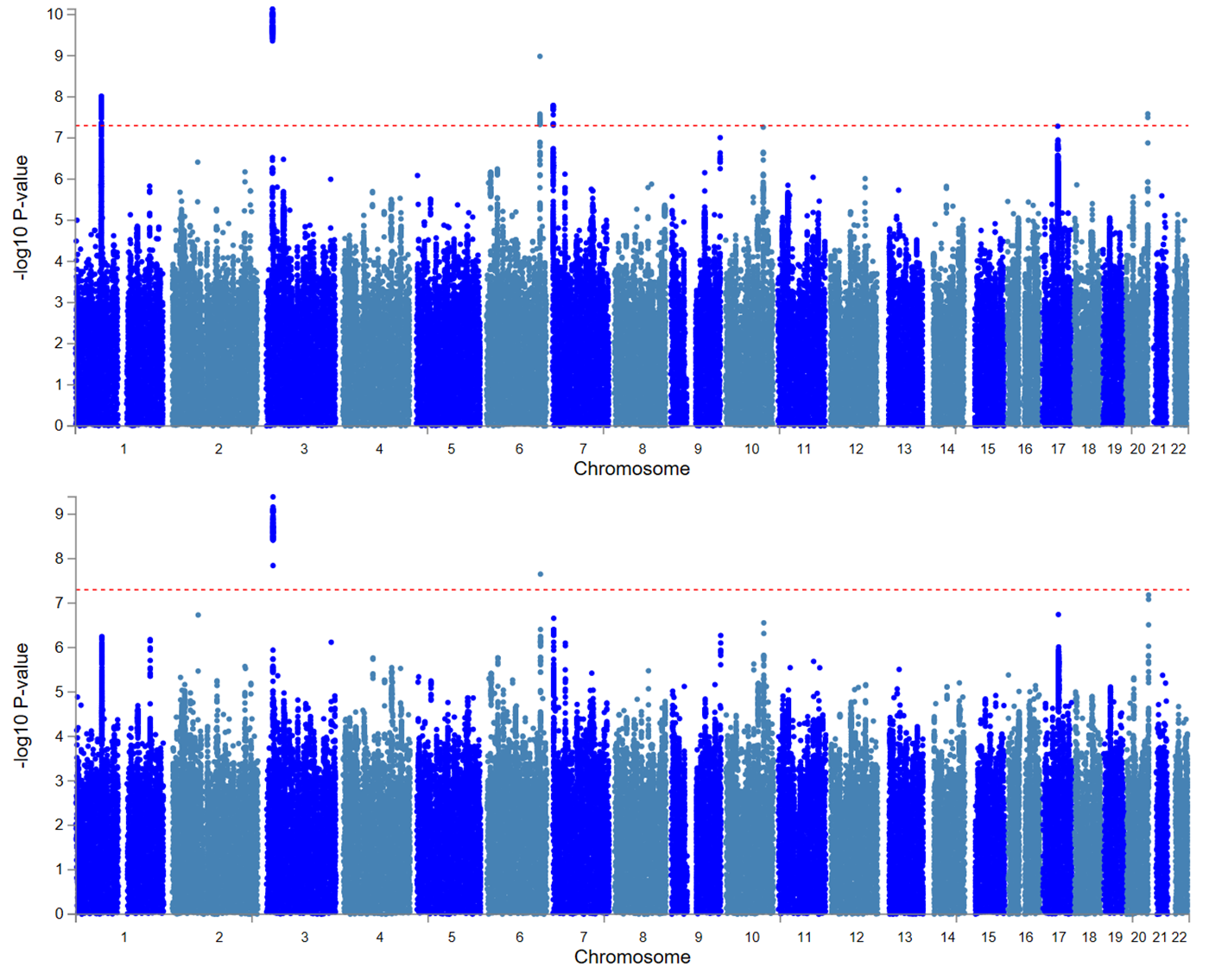

Supplementary Figure S4. GWGAS for European Americans

GAD-2

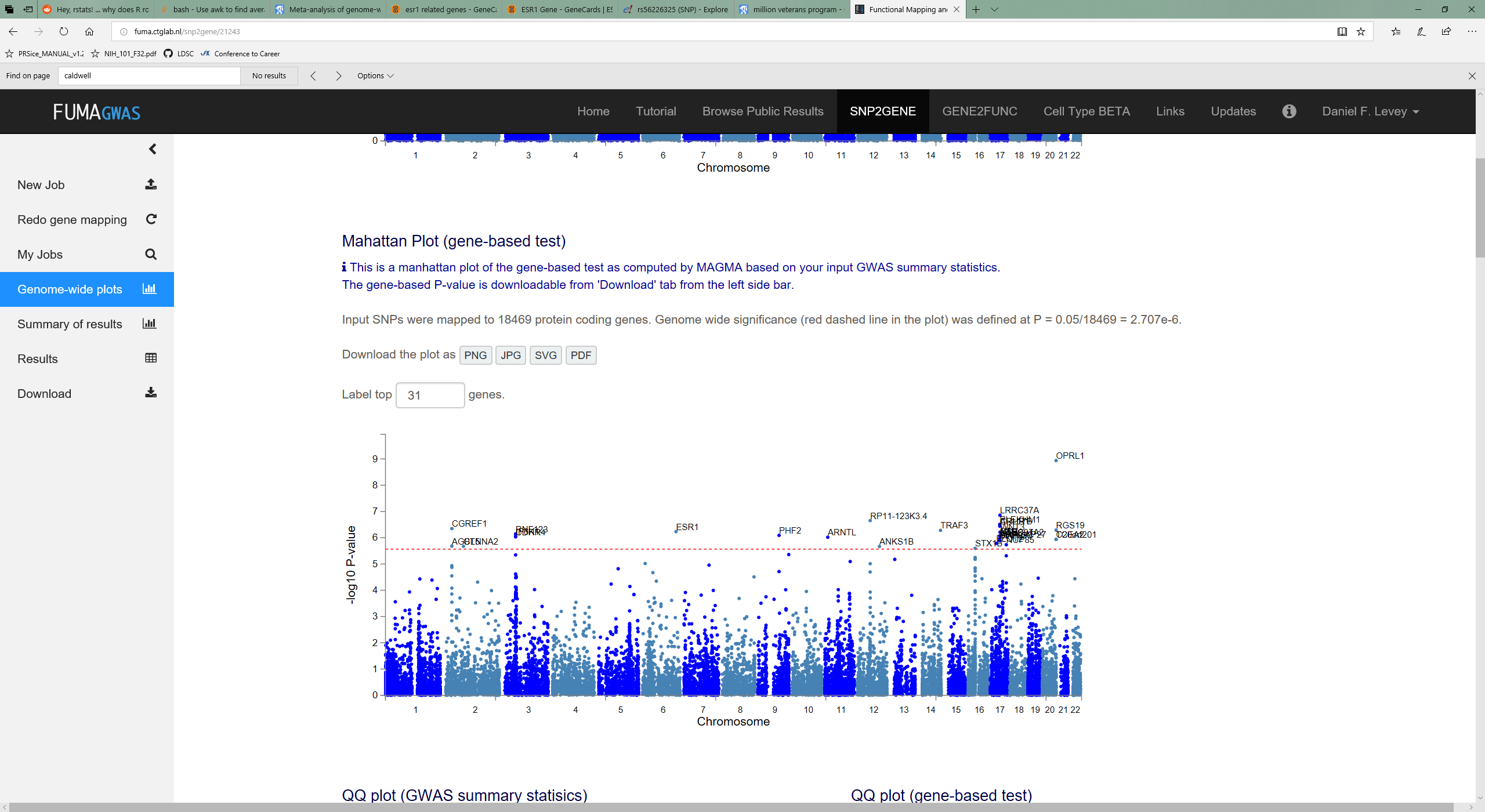

Case-Control

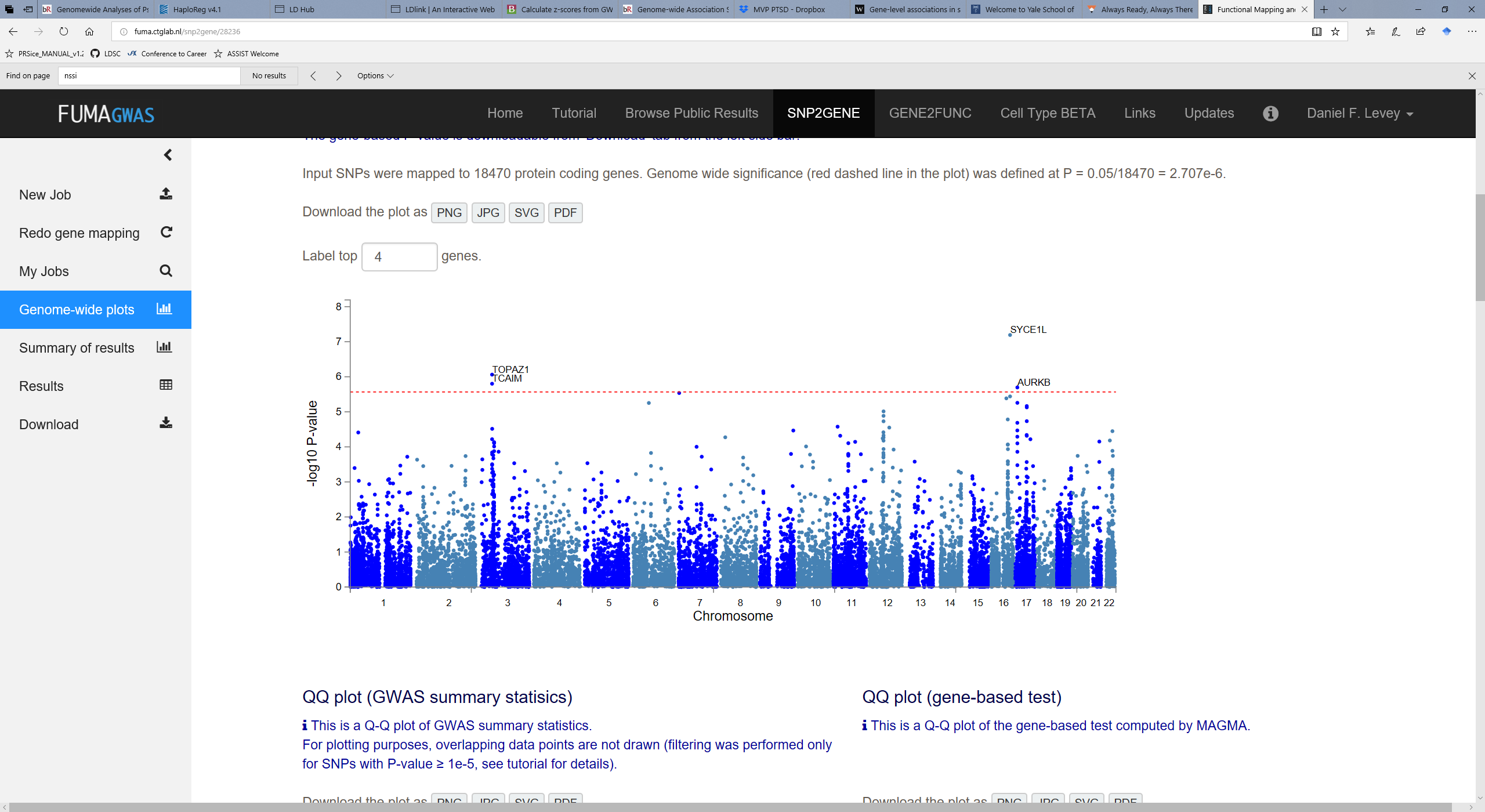

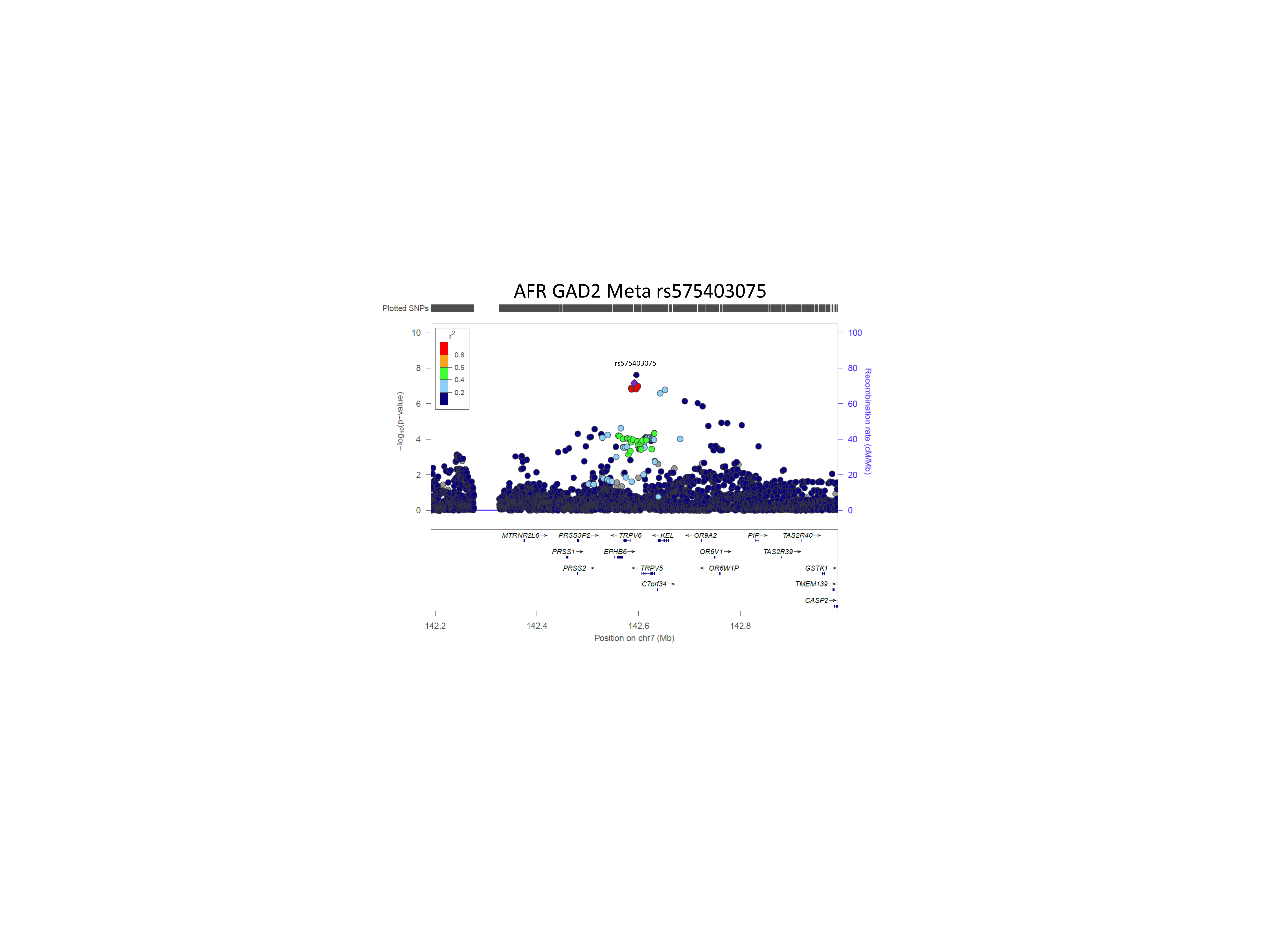

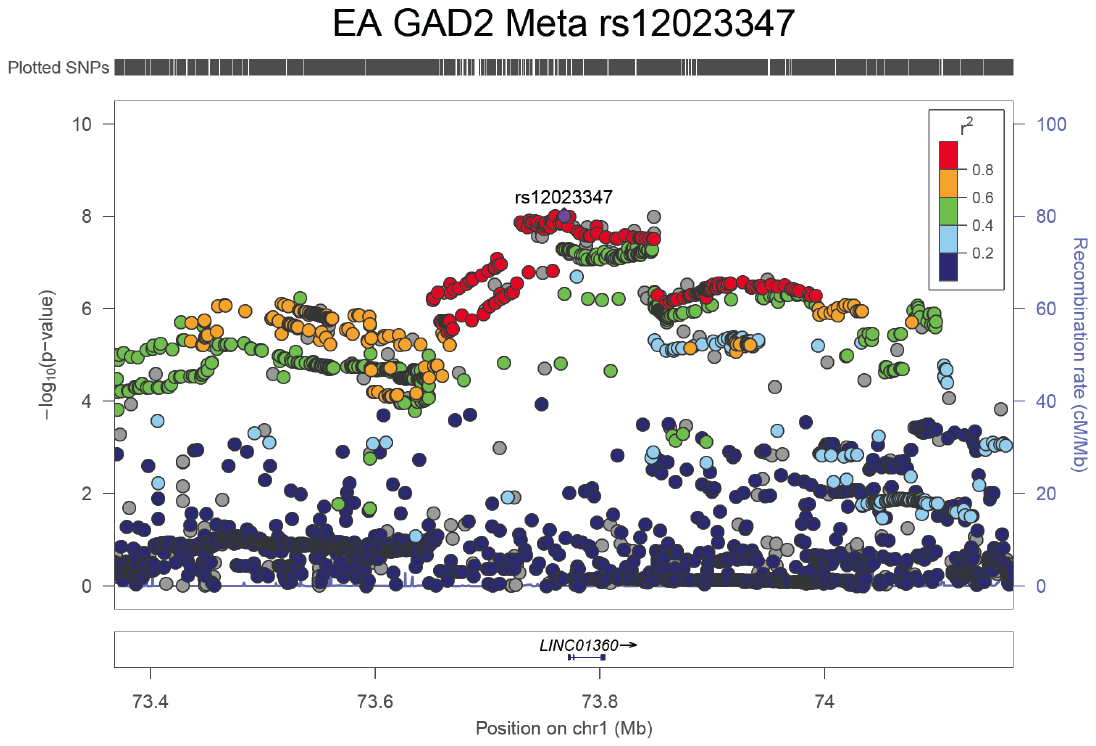

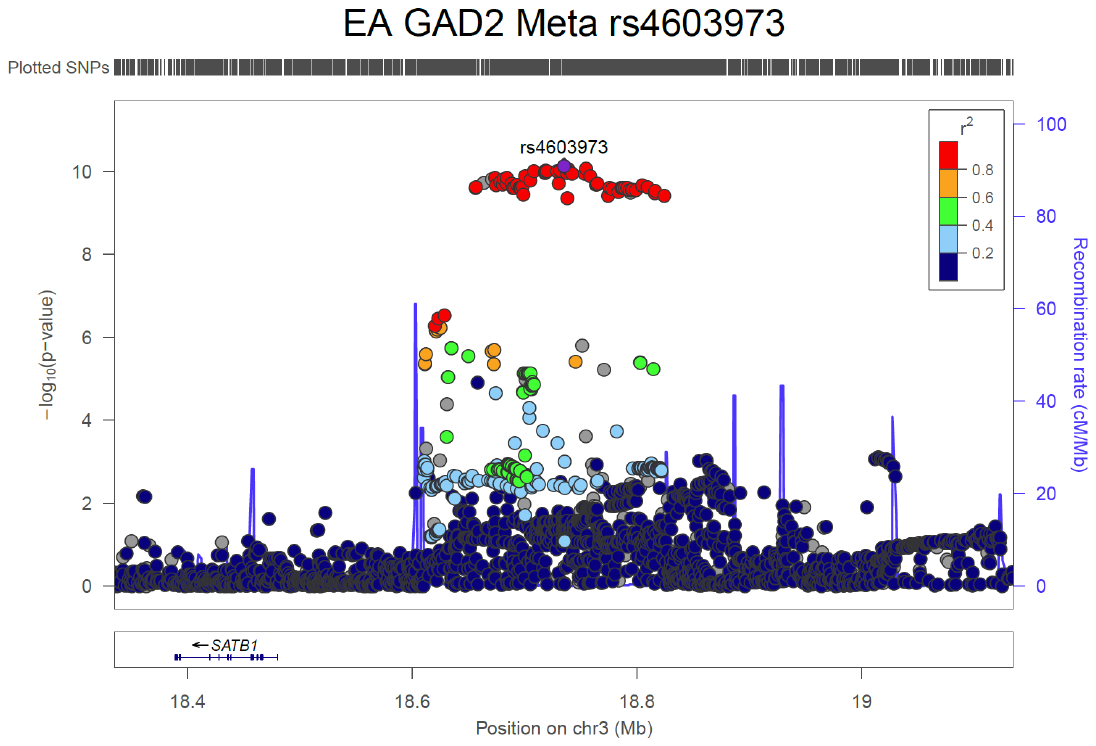

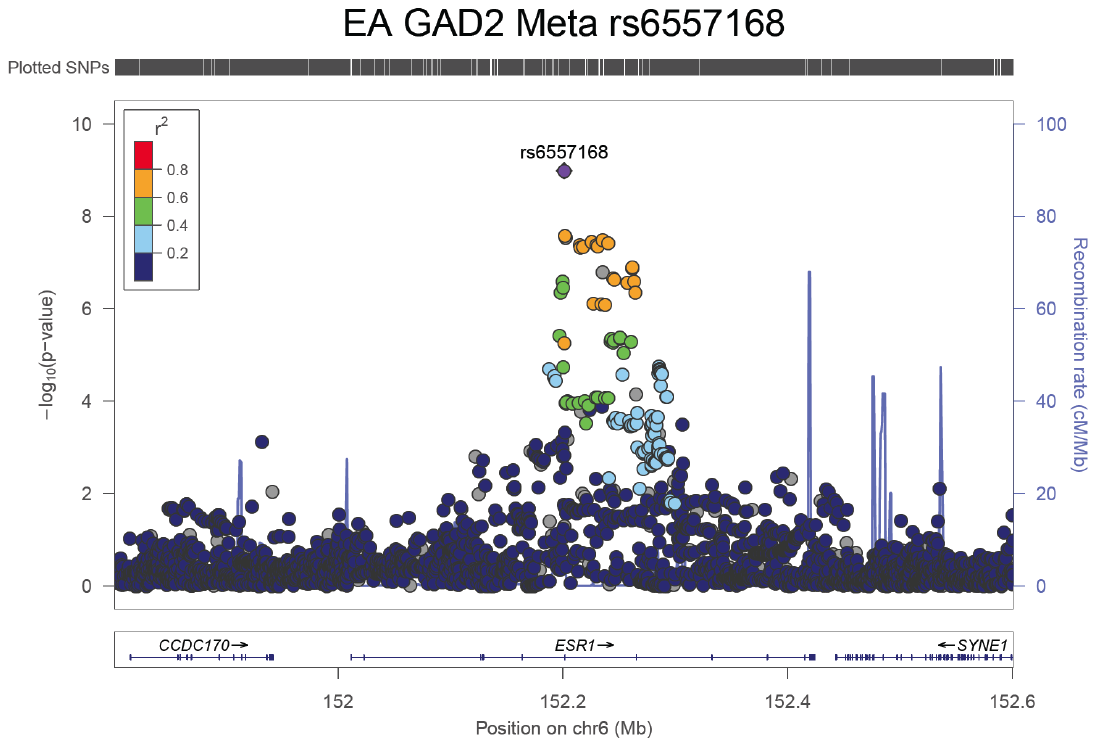

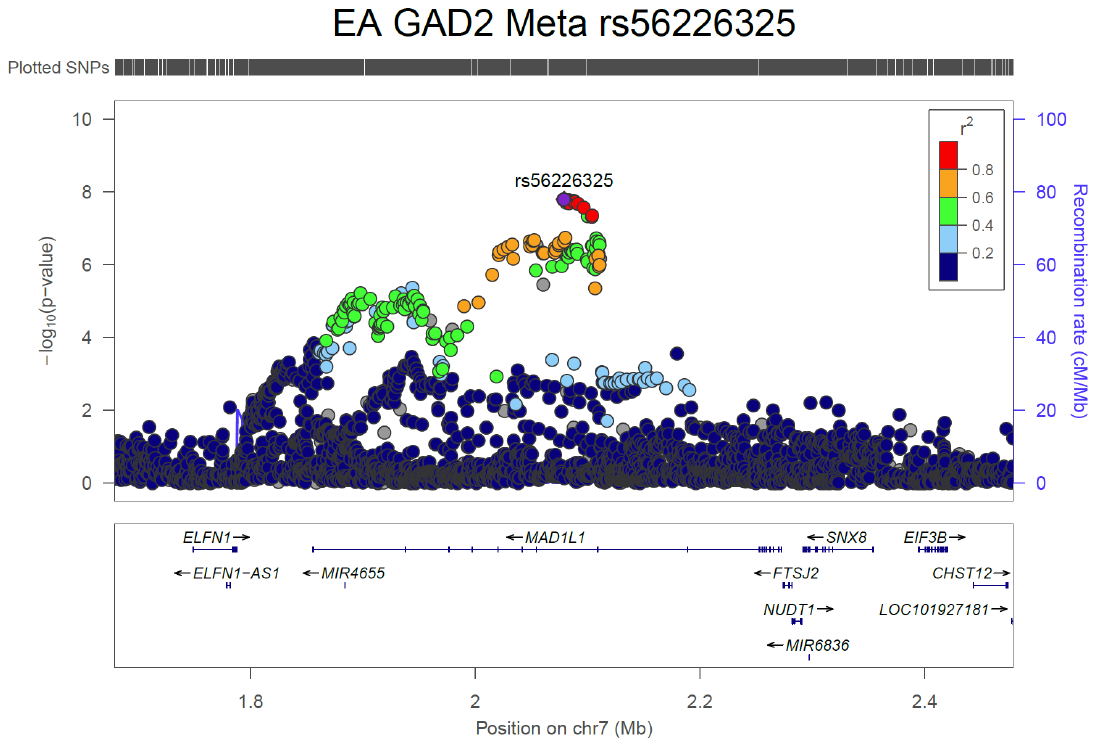

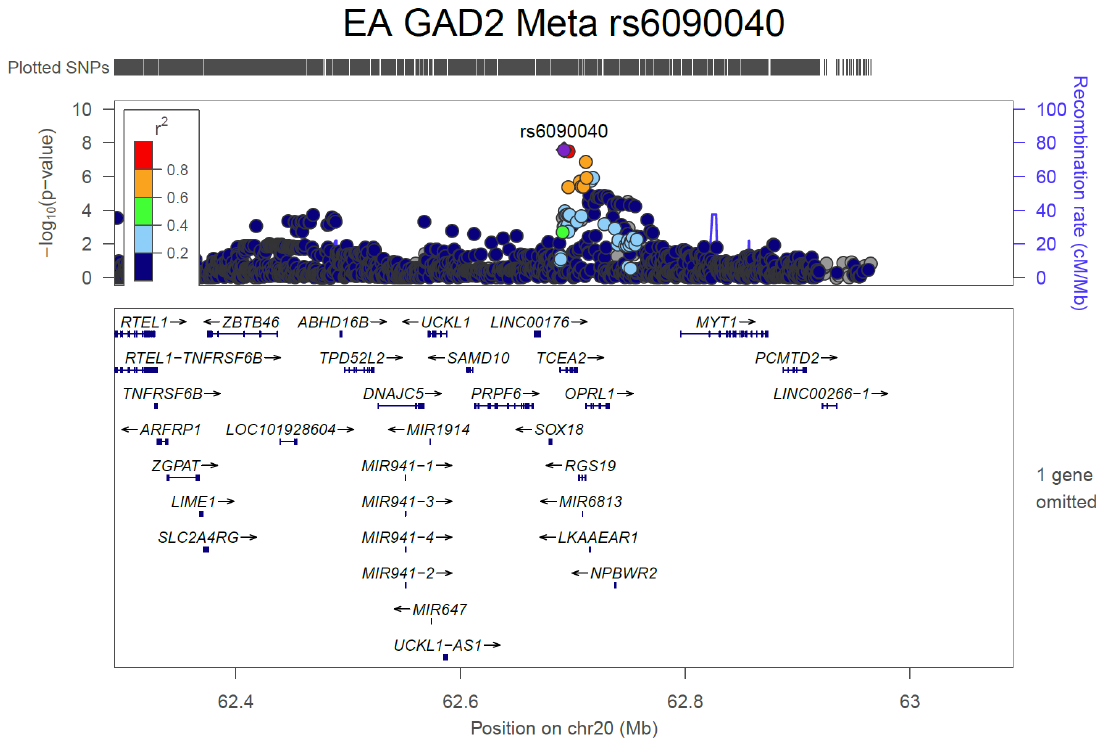

Supplementary Figure S5 Regional Manhattan Plots for GAD-2 GWAS.

Table S1 GAD-2 Phenotype

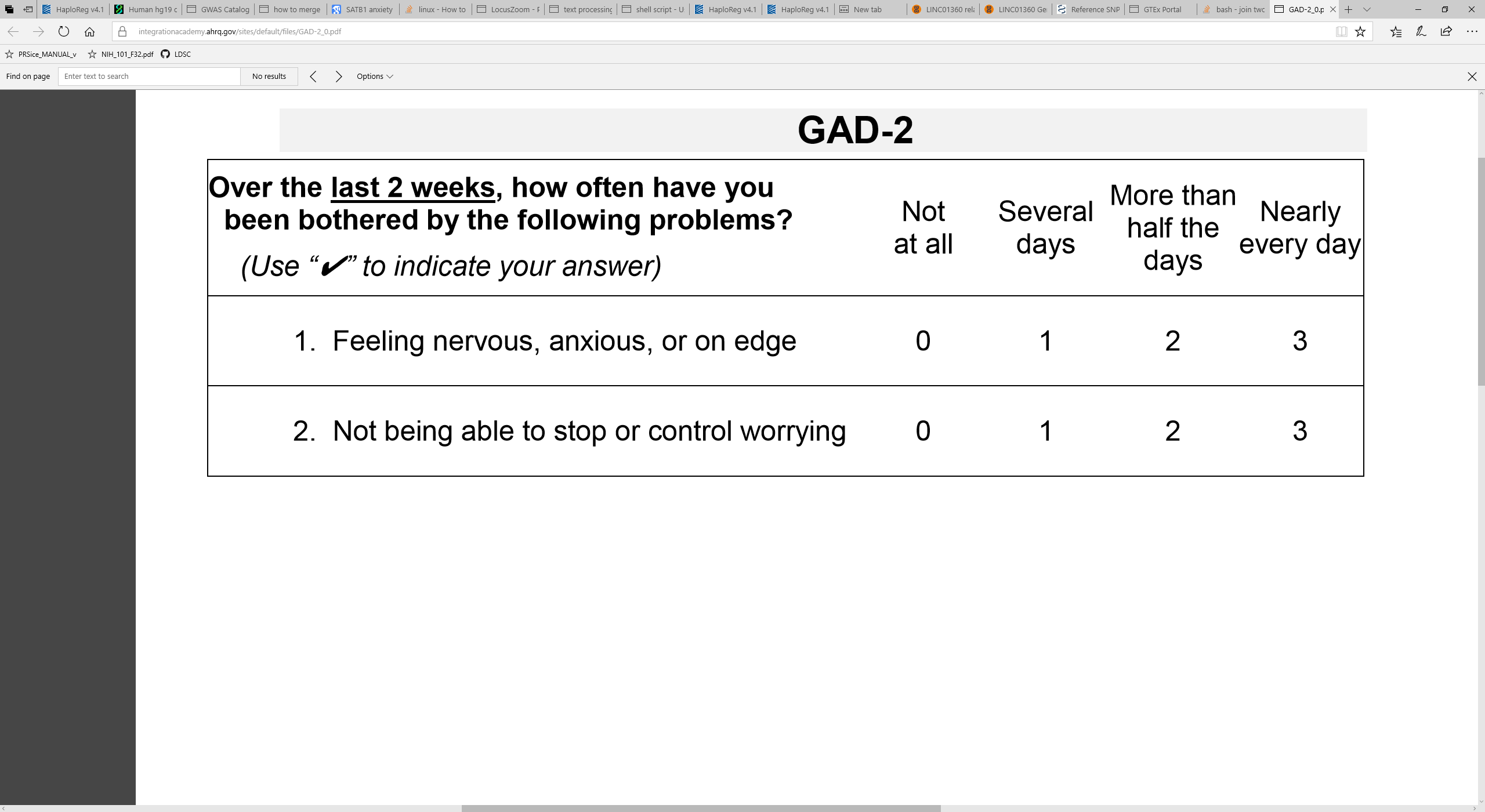

Table S2. Case control sample sizes

|  | Case | Control | Total |
| --- | --- | --- | --- |
| Self Reported Anx/Pan EUR | 28525 | 163731 | 192256 |
| Self Reported Anx/Pan AFR | 5664 | 26410 | 32074 |

Supplementary Table S3. Gene-Based Genome-Wide Association (GWGAS) in EAs.

| EA Meta GWGAS |  |  |  |  |  |  |  |  |  |
| --- | --- | --- | --- | --- | --- | --- | --- | --- | --- |
| GENE | CHR | START | STOP | NSNPS | NPARAM | N | ZSTAT | P | SYMBOL |
| ENSG00000125510 | 20 | 62711526 | 62731996 | 50 | 13 | 175163 | 5.9748 | 1.15E-09 | OPRL1 |
| ENSG00000176681 | 17 | 44370099 | 44415160 | 12 | 3 | 175163 | 5.1386 | 1.38E-07 | LRRC37A |
| ENSG00000120071 | 17 | 44107282 | 44302733 | 1014 | 6 | 175163 | 5.0888 | 1.80E-07 | KANSL1 |
| ENSG00000258830 | 12 | 57643392 | 57690267 | 72 | 14 | 175163 | 5.0489 | 2.22E-07 | RP11-123K3.4 |
| ENSG00000225190 | 17 | 43513266 | 43568115 | 157 | 8 | 175163 | 4.9858 | 3.09E-07 | PLEKHM1 |
| ENSG00000120088 | 17 | 43699267 | 43913194 | 1155 | 10 | 175163 | 4.9557 | 3.60E-07 | CRHR1 |
| ENSG00000228696 | 17 | 44352150 | 44439130 | 192 | 6 | 175163 | 4.9528 | 3.66E-07 | ARL17B |
| ENSG00000138028 | 2 | 27321757 | 27341995 | 47 | 9 | 175163 | 4.9128 | 4.49E-07 | CGREF1 |
| ENSG00000171700 | 20 | 62704534 | 62711323 | 12 | 5 | 175163 | 4.8871 | 5.12E-07 | RGS19 |
| ENSG00000131323 | 14 | 1.03E+08 | 1.03E+08 | 316 | 20 | 175163 | 4.8801 | 5.30E-07 | TRAF3 |
| ENSG00000091831 | 6 | 1.52E+08 | 1.52E+08 | 1673 | 89 | 175163 | 4.8585 | 5.91E-07 | ESR1 |
| ENSG00000108379 | 17 | 44839872 | 44910520 | 144 | 26 | 175163 | 4.8513 | 6.13E-07 | WNT3 |
| ENSG00000164068 | 3 | 49726932 | 49758962 | 52 | 9 | 175163 | 4.8205 | 7.16E-07 | RNF123 |
| ENSG00000197724 | 9 | 96338689 | 96441869 | 519 | 27 | 175163 | 4.79 | 8.34E-07 | PHF2 |
| ENSG00000256762 | 17 | 44076616 | 44077060 | 1 | 1 | 175163 | 4.7896 | 8.36E-07 | STH |
| ENSG00000073969 | 17 | 44668035 | 44834830 | 69 | 6 | 175163 | 4.7797 | 8.78E-07 | NSF |
| ENSG00000176095 | 3 | 49761727 | 49823975 | 131 | 11 | 175163 | 4.7768 | 8.91E-07 | IP6K1 |
| ENSG00000187492 | 3 | 49828165 | 49837268 | 18 | 7 | 175163 | 4.7686 | 9.28E-07 | CDHR4 |
| ENSG00000238083 | 17 | 44588877 | 44633016 | 14 | 4 | 175163 | 4.7666 | 9.37E-07 | LRRC37A2 |
| ENSG00000133794 | 11 | 13298199 | 13408813 | 307 | 37 | 175163 | 4.757 | 9.82E-07 | ARNTL |
| ENSG00000171695 | 20 | 62714733 | 62715712 | 3 | 1 | 175163 | 4.7232 | 1.16E-06 | C20orf201 |
| ENSG00000159314 | 17 | 43471275 | 43511787 | 115 | 8 | 175163 | 4.7217 | 1.17E-06 | ARHGAP27 |
| ENSG00000185294 | 17 | 43922256 | 43924438 | 17 | 2 | 175163 | 4.7139 | 1.22E-06 | SPPL2C |
| ENSG00000171703 | 20 | 62681189 | 62703700 | 72 | 17 | 175163 | 4.7125 | 1.22E-06 | TCEA2 |
| ENSG00000186868 | 17 | 43971748 | 44105700 | 822 | 5 | 175163 | 4.6821 | 1.42E-06 | MAPT |
| ENSG00000132589 | 17 | 27206353 | 27224697 | 30 | 6 | 175163 | 4.653 | 1.64E-06 | FLOT2 |
| ENSG00000125450 | 17 | 73201754 | 73231853 | 68 | 9 | 175163 | 4.6278 | 1.85E-06 | NUP85 |
| ENSG00000084693 | 2 | 27265232 | 27293490 | 46 | 9 | 175163 | 4.6021 | 2.09E-06 | AGBL5 |
| ENSG00000066032 | 2 | 79412357 | 80875905 | 4914 | 287 | 175163 | 4.5966 | 2.15E-06 | CTNNA2 |
| ENSG00000185046 | 12 | 99120235 | 1E+08 | 3499 | 186 | 175163 | 4.5933 | 2.18E-06 | ANKS1B |
| ENSG00000099365 | 16 | 31000577 | 31021949 | 47 | 10 | 175163 | 4.5538 | 2.63E-06 | STX1B |

Supplementary Table S4. Manually curated selection of top pathways from significant functional annotations (N=189 genes) using Ingenuity Pathway Analysis.

| Categories | Diseases or Functions Annotation | P-value | # Molecules |
| --- | --- | --- | --- |
| Cancer, Organismal Injury and Abnormalities | Carcinoma | 1.76E-07 | 156 |
| Cellular Assembly and Organization | Binding of membrane rafts | 6.20E-05 | 2 |
| Carbohydrate Metabolism, Small Molecule Biochemistry | Metabolism of glucose-1-phosphate | 6.20E-05 | 2 |
| Dermatological Diseases and Conditions, Organismal Injury and Abnormalities | Skin lesion | 1.21E-04 | 100 |
| Cancer, Dermatological Diseases and Conditions, Organismal Injury and Abnormalities | Skin tumor | 1.69E-04 | 99 |
| Inflammatory Response | Frequency of plasmacytoid dendritic cells | 1.85E-04 | 2 |
| Organismal Survival | Organismal death | 2.26E-04 | 48 |
| Behavior | Fear conditioning | 3.62E-04 | 3 |
| Gastrointestinal Disease, Hepatic System Disease, Organismal Injury and Abnormalities | Liver lesion | 8.84E-04 | 112 |
| Behavior, Reproductive System Development and Function | Mating | 7.80E-03 | 2 |

Supplementary Table S5

| Trait | PMID | Category | rg | se | z | p | h2_obs | h2_obs_se |
| --- | --- | --- | --- | --- | --- | --- | --- | --- |
| Depressive symptoms | 27089181 | psychiatric | 0.8055 | 0.0523 | 15.3885 | 1.95E-53 | 0.0482 | 0.0038 |
| Neuroticism1 | 27089181 | personality | 0.7174 | 0.0469 | 15.3103 | 6.53E-53 | 0.0904 | 0.0076 |
| Intelligence | 28530673 | cognitive | -0.4532 | 0.0415 | -10.9256 | 8.70E-28 | 0.1946 | 0.0104 |
| Age of first birth | 27798627 | reproductive | -0.4534 | 0.0444 | -10.2203 | 1.61E-24 | 0.0641 | 0.0036 |
| Subjective well being | 27089181 | psychiatric | -0.4511 | 0.0478 | -9.4447 | 3.56E-21 | 0.0251 | 0.0021 |
| Schizophrenia | 25056061 | psychiatric | 0.2687 | 0.0327 | 8.2183 | 2.06E-16 | 0.4601 | 0.019 |
| Insomnia1 | 27992416 | sleeping | 0.4395 | 0.0545 | 8.0662 | 7.25E-16 | 0.1323 | 0.0118 |
| Insomnia2 | 28604731 | sleeping | 0.4721 | 0.0646 | 7.3131 | 2.61E-13 | 0.0472 | 0.0048 |
| Neuroticism2 | 24828478 | personality | 0.7459 | 0.1086 | 6.8697 | 6.43E-12 | 0.0144 | 0.0034 |
| Major depressive disorder | 22472876 | psychiatric | 0.4768 | 0.0727 | 6.5554 | 5.55E-11 | 0.165 | 0.0259 |
| Number of children ever born | 27798627 | reproductive | 0.269 | 0.0477 | 5.6355 | 1.75E-08 | 0.0256 | 0.0018 |
| Ever vs never smoked | 20418890 | smoking behavior | 0.346 | 0.0624 | 5.5438 | 2.96E-08 | 0.0738 | 0.0067 |
| Former vs Current smoker | 20418890 | smoking_behavior | -0.4138 | 0.0871 | -4.75 | 2.03E-06 | 0.0602 | 0.0108 |
| PGC cross-disorder analysis | 23453885 | psychiatric | 0.2553 | 0.054 | 4.7305 | 2.24E-06 | 0.1711 | 0.0132 |
| Fathers age at death | 27015805 | aging | -0.3025 | 0.0731 | -4.1371 | 3.52E-05 | 0.0389 | 0.0066 |
| Mothers age at death | 27015805 | aging | -0.2967 | 0.0786 | -3.7735 | 0.0002 | 0.0351 | 0.0074 |
| Amyotrophic lateral sclerosis | 27455348 | neurological | 0.3852 | 0.1068 | 3.6065 | 0.0003 | 0.0452 | 0.0123 |
| Parents age at death | 27015805 | aging | -0.3041 | 0.0906 | -3.3561 | 0.0008 | 0.0292 | 0.007 |
| Attention deficit hyperactivity disorder (GC) | 27663945 | psychiatric | 0.382 | 0.1259 | 3.0339 | 0.0024 | 0.0698 | 0.0308 |
| Attention deficit hyperactivity disorder (No GC) | 27663945 | psychiatric | 0.381 | 0.126 | 3.0239 | 0.0025 | 0.0707 | 0.0313 |
| Parkinsons disease | 19915575 | neurological | -0.1908 | 0.0677 | -2.8207 | 0.0048 | 0.3724 | 0.1155 |
| Age at Menopause | 26414677 | reproductive | -0.1338 | 0.0501 | -2.6688 | 0.0076 | 0.1373 | 0.0169 |
| Anorexia Nervosa | 24514567 | psychiatric | 0.1056 | 0.0448 | 2.3568 | 0.0184 | 0.5495 | 0.0301 |
| Neo-openness to experience | 21173776 | personality | -0.2122 | 0.1012 | -2.0963 | 0.0361 | 0.1001 | 0.0267 |
| Bipolar disorder | 21926972 | psychiatric | 0.1028 | 0.0536 | 1.9172 | 0.0552 | 0.4445 | 0.0369 |
| Attention deficit hyperactivity disorder | 20732625 | psychiatric | 0.2345 | 0.1235 | 1.8993 | 0.0575 | 0.2366 | 0.0937 |
| Cigarettes smoked per day | 20418890 | smoking_behavior | 0.1544 | 0.0863 | 1.7884 | 0.0737 | 0.0543 | 0.016 |
| Chronotype | 27494321 | sleeping | 0.0633 | 0.0429 | 1.4758 | 0.14 | 0.1022 | 0.006 |
| Excessive daytime sleepiness | 27992416 | sleeping | 0.0718 | 0.0545 | 1.3179 | 0.1875 | 0.0533 | 0.0051 |
| Sleep duration | 27494321 | sleeping | -0.062 | 0.0518 | -1.1956 | 0.2318 | 0.057 | 0.0048 |
| Neo-conscientiousness | 21173776 | personality | -0.112 | 0.1154 | -0.9704 | 0.3319 | 0.075 | 0.029 |
| Age at Menarche | 25231870 | reproductive | -0.0268 | 0.0325 | -0.8222 | 0.411 | 0.2042 | 0.011 |
| Age of smoking initiation | 20418890 | smoking_behaviour | -0.0739 | 0.0953 | -0.7758 | 0.4379 | 0.0663 | 0.0194 |
| Alzheimers disease | 24162737 | neurological | 0.0005 | 0.0841 | 0.0065 | 0.9948 | 0.0459 | 0.0224 |

Supplementary Table S6. Genetic overlap between GAD-2 trait anxiety and Anxiety/Panic disorder or treatment received for Anxiety/Panic disorder.

| trait | rg | se | z | p | h2_obs | h2_obs_se |
| --- | --- | --- | --- | --- | --- | --- |
| Anxiety/Panic Disorder | 0.8557 | 0.0368 | 23.2277 | 2.39E-119 | 0.0374 | 0.0037 |
| Treatment for Anxiety/Panic | 0.7962 | 0.0444 | 17.9398 | 5.77E-72 | 0.0313 | 0.0034 |

Supplementary Table S7. eQTL evidence for top GWS SNPs.

| uniqID | db | tissue | gene | testedAllele | p | signed_stats | FDR | RiskIncAllele | chr | pos | symbol | eqtlMapFilt |
| --- | --- | --- | --- | --- | --- | --- | --- | --- | --- | --- | --- | --- |
| 20:62706105:A:G | GTEx_v7 | Brain_Caudate_basal_ganglia | ENSG00000171695 | G | 9.40E-06 | -0.357702 | 0.00916956 | G | 20 | 62706105 | C20orf201 | 1 |
| 20:62707527:C:T | GTEx_v7 | Brain_Caudate_basal_ganglia | ENSG00000171695 | T | 1.07E-05 | -0.363151 | 0.00916956 | T | 20 | 62707527 | C20orf201 | 1 |
| 20:62709274:A:G | GTEx_v7 | Brain_Caudate_basal_ganglia | ENSG00000171695 | A | 2.43E-06 | -0.380321 | 0.00916956 | A | 20 | 62709274 | C20orf201 | 1 |
| 20:62712053:C:T | GTEx_v7 | Brain_Caudate_basal_ganglia | ENSG00000171695 | C | 3.95E-06 | -0.368909 | 0.00916956 | C | 20 | 62712053 | C20orf201 | 1 |
| 7:2085165:C:T | GTEx_v7 | Brain_Cerebellar_Hemisphere | ENSG00000122687 | C | 6.28E-06 | -0.297022 | 8.17E-08 | T | 7 | 2085165 | FTSJ2 | 1 |
| 7:2085553:C:T | GTEx_v7 | Brain_Cerebellar_Hemisphere | ENSG00000122687 | C | 6.28E-06 | -0.297022 | 8.17E-08 | T | 7 | 2085553 | FTSJ2 | 1 |
| 20:62692060:A:C | GTEx_v7 | Brain_Cerebellar_Hemisphere | ENSG00000171700 | C | 9.75E-07 | 0.452831 | 0.00257718 | C | 20 | 62692060 | RGS19 | 1 |
| 20:62695931:A:G | GTEx_v7 | Brain_Cerebellar_Hemisphere | ENSG00000125510 | A | 2.53E-07 | 0.444876 | 0.000539622 | A | 20 | 62695931 | OPRL1 | 1 |
| 20:62695931:A:G | GTEx_v7 | Brain_Cerebellar_Hemisphere | ENSG00000171700 | A | 2.19E-06 | 0.429128 | 0.00257718 | A | 20 | 62695931 | RGS19 | 1 |
| 20:62696024:C:T | GTEx_v7 | Brain_Cerebellar_Hemisphere | ENSG00000171700 | T | 6.24E-06 | 0.424968 | 0.00257718 | T | 20 | 62696024 | RGS19 | 1 |
| 20:62706105:A:G | GTEx_v7 | Brain_Cerebellar_Hemisphere | ENSG00000125510 | G | 7.56E-08 | 0.457153 | 0.000539622 | G | 20 | 62706105 | OPRL1 | 1 |
| 20:62706105:A:G | GTEx_v7 | Brain_Cerebellar_Hemisphere | ENSG00000171700 | G | 5.36E-06 | 0.410154 | 0.00257718 | G | 20 | 62706105 | RGS19 | 1 |
| 20:62707527:C:T | GTEx_v7 | Brain_Cerebellar_Hemisphere | ENSG00000125510 | T | 3.04E-07 | 0.439947 | 0.000539622 | T | 20 | 62707527 | OPRL1 | 1 |
| 20:62707527:C:T | GTEx_v7 | Brain_Cerebellar_Hemisphere | ENSG00000171700 | T | 8.12E-06 | 0.404679 | 0.00257718 | T | 20 | 62707527 | RGS19 | 1 |
| 20:62709274:A:G | GTEx_v7 | Brain_Cerebellar_Hemisphere | ENSG00000125510 | A | 8.76E-08 | 0.449324 | 0.000539622 | A | 20 | 62709274 | OPRL1 | 1 |
| 20:62709274:A:G | GTEx_v7 | Brain_Cerebellar_Hemisphere | ENSG00000171700 | A | 2.37E-06 | 0.418317 | 0.00257718 | A | 20 | 62709274 | RGS19 | 1 |
| 20:62712053:C:T | GTEx_v7 | Brain_Cerebellar_Hemisphere | ENSG00000125510 | C | 6.82E-08 | 0.446404 | 0.000539622 | C | 20 | 62712053 | OPRL1 | 1 |
| 20:62712053:C:T | GTEx_v7 | Brain_Cerebellar_Hemisphere | ENSG00000171700 | C | 1.46E-06 | 0.420188 | 0.00257718 | C | 20 | 62712053 | RGS19 | 1 |
| 20:62692060:A:C | GTEx_v7 | Brain_Cerebellum | ENSG00000171700 | C | 9.42E-08 | 0.435066 | 0.000132103 | C | 20 | 62692060 | RGS19 | 1 |
| 20:62695931:A:G | GTEx_v7 | Brain_Cerebellum | ENSG00000125510 | A | 5.26E-07 | 0.388893 | 0.000621703 | A | 20 | 62695931 | OPRL1 | 1 |
| 20:62695931:A:G | GTEx_v7 | Brain_Cerebellum | ENSG00000171700 | A | 1.88E-08 | 0.446254 | 0.000132103 | A | 20 | 62695931 | RGS19 | 1 |
| 20:62696024:C:T | GTEx_v7 | Brain_Cerebellum | ENSG00000125510 | T | 3.42E-05 | 0.333873 | 0.000621703 | T | 20 | 62696024 | OPRL1 | 1 |
| 20:62696024:C:T | GTEx_v7 | Brain_Cerebellum | ENSG00000171700 | T | 6.54E-08 | 0.440536 | 0.000132103 | T | 20 | 62696024 | RGS19 | 1 |
| 20:62706105:A:G | GTEx_v7 | Brain_Cerebellum | ENSG00000125510 | G | 1.10E-07 | 0.411089 | 0.000621703 | G | 20 | 62706105 | OPRL1 | 1 |
| 20:62706105:A:G | GTEx_v7 | Brain_Cerebellum | ENSG00000171700 | G | 5.23E-08 | 0.435991 | 0.000132103 | G | 20 | 62706105 | RGS19 | 1 |
| 20:62707527:C:T | GTEx_v7 | Brain_Cerebellum | ENSG00000125510 | T | 1.55E-07 | 0.413277 | 0.000621703 | T | 20 | 62707527 | OPRL1 | 1 |
| 20:62707527:C:T | GTEx_v7 | Brain_Cerebellum | ENSG00000171700 | T | 1.08E-07 | 0.433475 | 0.000132103 | T | 20 | 62707527 | RGS19 | 1 |
| 20:62709274:A:G | GTEx_v7 | Brain_Cerebellum | ENSG00000125510 | A | 1.04E-07 | 0.410003 | 0.000621703 | A | 20 | 62709274 | OPRL1 | 1 |
| 20:62709274:A:G | GTEx_v7 | Brain_Cerebellum | ENSG00000171700 | A | 6.10E-08 | 0.432145 | 0.000132103 | A | 20 | 62709274 | RGS19 | 1 |
| 20:62712053:C:T | GTEx_v7 | Brain_Cerebellum | ENSG00000125510 | C | 9.95E-08 | 0.408897 | 0.000621703 | C | 20 | 62712053 | OPRL1 | 1 |
| 20:62712053:C:T | GTEx_v7 | Brain_Cerebellum | ENSG00000171700 | C | 1.28E-07 | 0.420938 | 0.000132103 | C | 20 | 62712053 | RGS19 | 1 |
